# Expressed barcodes enable clonal characterization of chemotherapeutic responses in chronic lymphocytic leukemia

**DOI:** 10.1101/761981

**Authors:** Aziz Al’Khafaji, Catherine Gutierrez, Eric Brenner, Russell Durrett, Kaitlyn E. Johnson, Wandi Zhang, Shuqiang Li, Kenneth J. Livak, Donna Neuberg, Amy Brock, Catherine J. Wu

## Abstract

The remarkable evolutionary capacity of cancer is a major challenge to current therapeutic efforts. Fueling this evolution is its vast clonal heterogeneity and ability to adapt to diverse selective pressures. Although the genetic and transcriptional mechanisms underlying these responses have been independently evaluated, the ability to couple genetic alterations present within individual clones to their respective transcriptional or functional outputs has been lacking in the field. To this end, we developed a high-complexity expressed barcode library that integrates DNA barcoding with single-cell RNA sequencing through use of the CROP-seq sgRNA expression/capture system, and which is compatible with the COLBERT clonal isolation workflow for subsequent genomic and epigenomic characterization of specific clones of interest. We applied this approach to study chronic lymphocytic leukemia (CLL), a mature B cell malignancy notable for its genetic and transcriptomic heterogeneity and variable disease course. Here, we demonstrate the clonal composition and gene expression states of HG3, a CLL cell line harboring the common alteration *del*(13q), in response to front-line cytotoxic therapy of fludarabine and mafosfamide (an analog of the clinically used cyclophosphamide). Analysis of clonal abundance and clonally-resolved single-cell RNA sequencing revealed that only a small fraction of clones consistently survived therapy. These rare highly drug tolerant clones comprise 94% of the post-treatment population and share a stable, pre-existing gene expression state characterized by upregulation of CXCR4 and WNT signaling and a number of DNA damage and cell survival genes. Taken together, these data demonstrate at unprecedented resolution the diverse clonal characteristics and therapeutic responses of a heterogeneous cancer cell population. Further, this approach provides a template for the high-resolution study of thousands of clones and the respective gene expression states underlying their response to therapy.

## Introduction

Tumors are composed of a heterogeneous population of cells with variable genetic and epigenetic characteristics that together contribute to their ability to adapt and further diversify in response to their environment. It is precisely this intrinsic heterogeneity that fuels cancer’s powerful evolutionary capacity and highlights why many therapeutic successes are so short-lived. The advent of high-throughput sequencing efforts has enabled deeper characterizations of the genomic and transcriptomic profiles that arise as a result of evolution, such that the field has gained a great appreciation for the diverse ways by which cancer cells can evade therapy. The rational development of treatment that yields durable responses will require a more granular understanding of the processes by which individual cells arrive at each of their successful evolutionary outcomes.

Clonal evolution of cancer has been extensively studied by tracking variant allele frequencies over time using whole genome or whole exome sequencing (WGS/WES) (Egan et al., 2012; Kasar et al., 2015; Landau et al., 2013, 2015; Roth et al., 2014), yet this approach is inference-based and currently limited by the lower bounds of sequencing detection. Recent efforts have increasingly leveraged DNA barcoding, a lineage tracing approach that introduces a unique, heritable marker into the genome of individual cells such that they and their descendants can be tagged and tracked over time (Bhang et al., 2015; Kalhor et al., 2017, 2018; McKenna et al., 2016; Pei et al., 2017). Using this method, previous groups have identified the presence of pre-existing and *de novo* mutations that can respectively drive resistance to therapy (Bhang et al., 2015; Hata et al., 2016).

While these approaches can highlight changes in clonal abundances over time, they are limited in their ability to integrate genomic, epigenomic and transcriptomic signatures with clonal identity. To address this challenge, we developed a novel methodology that combines traditional DNA barcoding with CROP-seq, a single guide-RNA (sgRNA) expression vector that implements the expression and capture of barcodes in single-cell RNA sequencing (scRNA-seq) workflows (Datlinger et al., 2017). In doing so, we couple the strengths of lineage tracing with the high-resolution study of individual cells through analyses such as scRNA-seq. Furthermore, this barcoded sgRNA system is compatible with the COLBERT workflow for clonal isolation, which enables downstream clone characterization (of genomic features, such as by WGS/WES, or epigenomic features, such as by ATAC-seq) and live-cell manipulation of these complex populations (Al’Khafaji et al., 2018).

We have applied this combined methodology to study the nature of clonal responses to chemotherapy in a chronic lymphocytic leukemia (CLL) cell line, HG3, previously described to be genetically heterogeneous in composition (Quentmeier et al., 2016). CLL is a typically indolent B cell malignancy with a diverse clinical course across patients. Like many other cancers, it has been characterized through several large-scale sequencing efforts as having a high degree of inter- and intra-tumoral heterogeneity, with no one mutation solely responsible for driving disease progression or therapeutic resistance. Though CLL’s therapeutic landscape has evolved considerably over the last 5 years, the longest experience and clinical outcome data is associated with frontline treatment with chemotherapy combinations that include fludarabine and cyclophosphamide. Novel targeted agents such as inhibitors of BTK and BCL2 signaling (i.e. ibrutinib and venetoclax) have emerged as promising therapies, yet relapse to these agents, as in the case of chemo-immunotherapies, remains a major challenge. Understanding the varied evolutionary paths and processes that enable cancer cell populations to evade therapy will thus greatly impact our ability to predict disease prognosis, rate of progression and likelihood of relapse, as well as potentially identify novel targets for therapy. Here, we investigate these properties, using transcriptome-wide sequencing coupled with lineage tracing methodologies to elucidate the mechanisms underlying clonal dynamics and resistance to frontline chemotherapy in HG3 cells.

## Results

To uncover the heterogeneity of HG3 cells while also characterizing the genomic and transcriptomic profiles of respective clones, we constructed a highly diverse DNA barcode library using a CROP-seq variant that expresses blue fluorescent protein (BFP) as a reporter with fully degenerate N(20) barcodes integrating into the 3’ UTR sgRNA cassette (**Fig. 1a-b**). Deep sequencing of the assembled barcode library vector confirmed its high diversity and even distribution; by fitting the sequencing depth to the number of barcodes observed through NGS, we projected that the vector library contained ~7.6×10^7^ unique barcodes (**Fig. 1c**) and identified ~6.8×10^7^ unique barcodes with 99.95% of barcodes detected falling below 10 reads (**Fig. 1d**).

**Figure 1:**
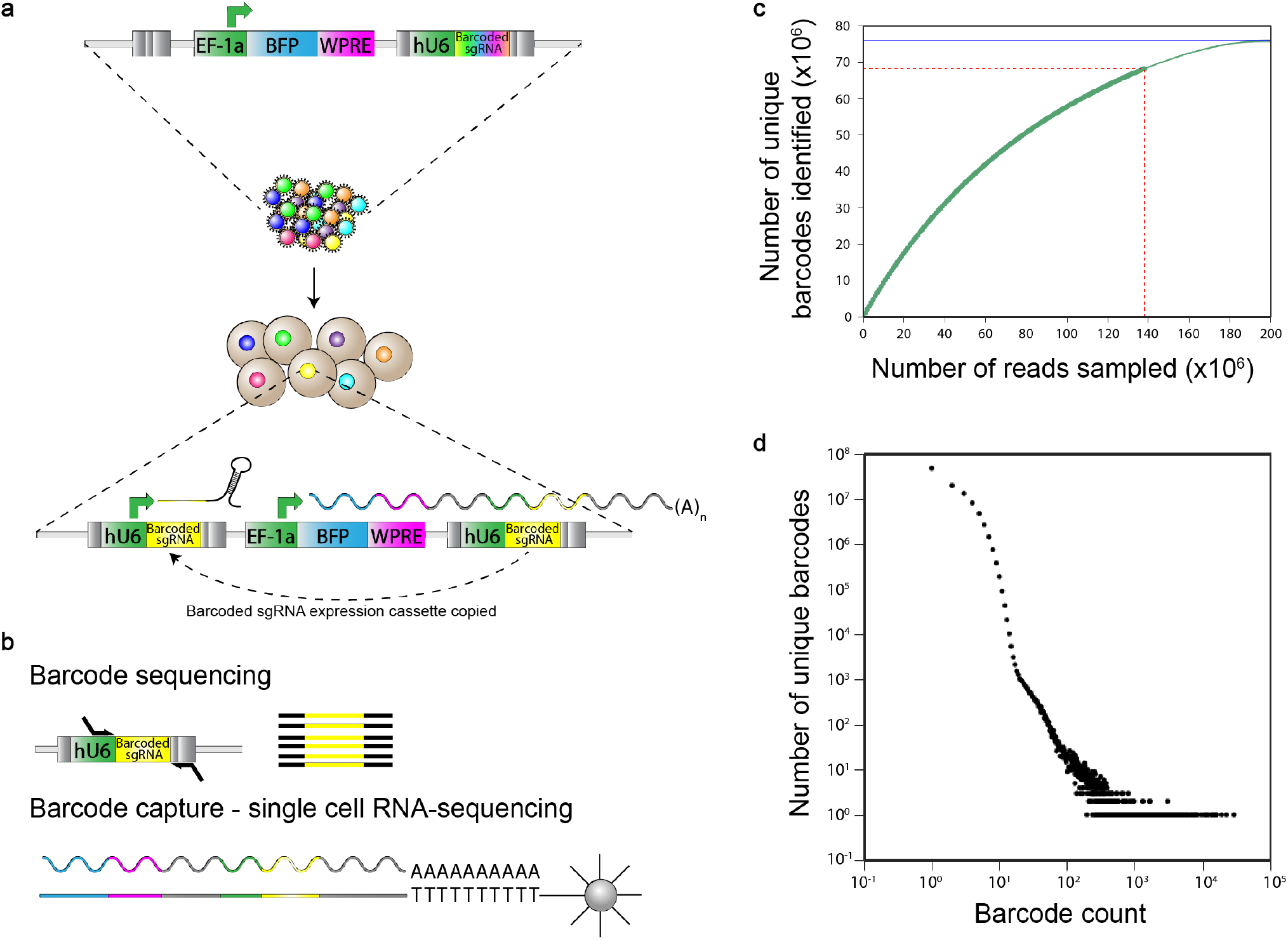
Generation of the expressed barcode-sgRNA library. (a) The CROP-seq lentiviral vector was modified to express BFP as well as a library of barcode-sgRNAs. The expressed barcode-sgRNA cassette is located in the 3’UTR and is duplicated upon genomic integration. (b) This expressed-barcode sgRNA construct generates a heritable DNA tag quantifiable by deep sequencing, expression of a CRISPR compatible barcode-sgRNA, and a poly-adenylated barcode transcript that can be captured in scRNA-seq workflows. (c) Barcode library diversity of the CROP-seq barcode-sgRNA library was estimated by deep sequencing. The dark green points correspond to the total unique read counts at corresponding read depths. Approximately 68 million unique barcodes were identified at a read depth of 138 million reads, indicated by the dashed red line. By fitting the sampled data, denoted by the blue line, the total number of unique barcodes is estimated to be approximately 7.6×10^6^. (d) The barcode frequency distribution of the CROP-seq barcode-sgRNA library was similarly quantified by deep sequencing at a depth of 138 million reads with no base below Q30. No significant bias was observed, with 99.95% of the 68 million unique barcodes present ≤10 times, with the most abundant barcode present in only 20,857 reads (0.03% of the population).

We transduced this barcode library into the CLL cell line HG3 at a multiplicity of infection (MOI) of 0.1 to minimize multiple barcode integrations into any given cell. Approximately 1.2×10^6^ BFP^+^ HG3 cells were collected and expanded for 27 days (~13 doublings) to establish the parental barcoded HG3 population (**Fig 2a**) and were subsequently aliquoted and cryopreserved. To study the response of this barcoded HG3 population to the frontline combination chemotherapy regimen of fludarabine and cyclophosphamide (using the active *in vitro* analog mafosfamide in lieu of cyclophosphamide), a vial containing 15 million live cells was thawed, expanded for 8 days (~4 doublings) and divided into 8 parallel replicates of 1×10^7^ cells each, ensuring a comparable barcode starting representation across replicates. These parallel replicates were then treated with an LD95 combined dose of fludarabine (5 μM) and mafosfamide (5 μM) for 72 hours to generate a deep population bottleneck. The cells were thereafter washed and resuspended in fresh media for outgrowth. As in previous studies, we reasoned that clones consistently enriched by treatment across replicates likely harbor a pre-existing mechanism of drug tolerance (Bhang et al., 2015).

**Figure 2:**
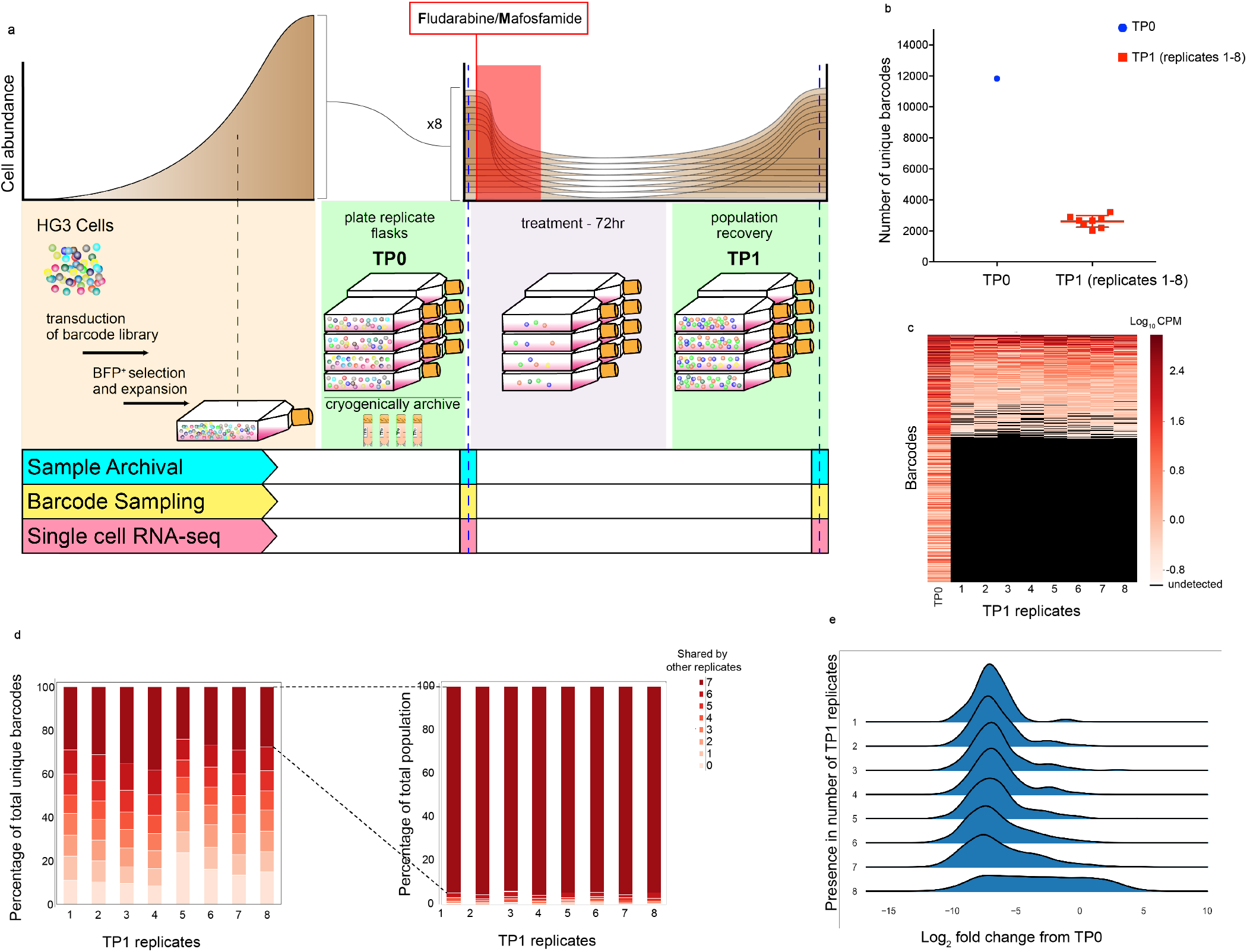
Measurement of clonal dynamics in response to treatment. (a) Experimental workflow for parallel treatment of barcoded HG3 population. HG3 cells transduced with the lentiviral barcode library at a MOI of 0.1 were isolated by FACS, expanded and either cryopreserved or plated in 8 parallel replicates. Replicates were treated with an LD95 combined dose of fludarabine (5 μM) and mafosfamide (5 μM) for 72 hours, washed and monitored for 21 days until outgrowth. Pre- and post-treatment samples were either processed for barcode sampling or cryopreserved for later scRNA-sequencing analysis. (b) Barcode sampling of TP0 (in blue) and TP1 replicates 1-8 (in red) demonstrate a marked decrease in barcode diversity (11,827 to 2,622 ± 380, n=8; or ~78%). (c) Heatmap of unique barcode read counts at TP0 and TP1 replicates 1-7 normalized to log10 counts per million. Barcodes are sorted in descending order by the sum of their counts across the TP1 columns. (d) Stacked bar graph showing the percentage of barcodes in each TP1 replicate also found in 1 or more other replicates. Shades of red denote the number of additional replicates in which the barcodes were identified. Approximately 68.24% ± 5.76% of barcodes were found in 4 or more replicates, with 30.2% ± 4.62% found in all 8 replicates. Second stacked bar graph demonstrating that overall, the 30% of barcodes found in all replicates comprised 94.47% ±0.62% of the total barcode count in each replicate. (e) Ridgeline plot representing the log2 fold change of barcodes from TP0 to TP1, where barcodes are grouped by their presence across replicates. Only barcodes at or above a 0.01% abundance at TP0 are shown. Barcodes with a log2 fold change above 0 tend to be found in all 8 replicates (1082/1166).

We observed a massive decrease in viability across all 8 replicates, with eventual population recovery 20 days after fludarabine and mafosfamide treatment. Genomic DNA was isolated after minimal cell expansion and survivor populations were cryopreserved for later investigations. Deep sequencing of both pre-treatment (TP0) and fludarabine/mafosfamide recovered (TP1) populations revealed an approximately 80% decrease in the number of detectable unique clones (**Fig 2b**). Additionally, we found the clonal composition to be heavily skewed following treatment, with a small number of clones comprising the majority of the TP1 populations (**Fig 2c**). Along with similar trends in time to recovery and clonal diversity, we discovered that ~30% of the unique clones present in one TP1 replicate were shared by all other replicates (**Fig 2d**). Furthermore, we observed that the vast majority of clones that increased in abundance (i.e. a positive log2fold change from TP0) were present in all 8 replicates and totaled ~94% of the composite populations – indicating the likely presence of pre-existing mechanisms underlying survival (**Fig 2d-inset, Fig 2e**).

This clonal consistency in survival indicated the potential presence of a pre-existing drug tolerant state providing the phenotypic underpinnings of this differential survival. To this end, we performed scRNA-seq on 2 of 8 replicates collected at TP1 (replicates 1 and 7) and a pretreatment sample from TP0 using the 10x Chromium single-cell platform. We used Uniform Manifold Approximation and Projection (UMAP), a dimensional reduction graphical representation, to visualize the single-cell gene expression data (Becht et al., 2019). We found that each sample (TP0 and TP1, replicates 1 and 7) broadly segregated by treatment status and were comprised of multiple distinct clusters (**Fig 3a**). To identify clusters by gene expression, we constructed nearest neighbors graphs with clusters assigned (Leiden algorithm, see Methods). Leiden clustering resulted in assignment of 2 clusters associated with TP0 and 6 clusters comprising TP1 (**Fig 3b**). As our scRNA-seq data contained captured expressed barcodes, we could annotate the transcriptionally defined clusters within the UMAP with the 10 most abundant clones at TP1 (“top 10 clones”). This revealed that all but one cluster (cluster 6) at TP1 contained a top 10 clone (**Fig 3c**). Furthermore, when focusing on TP0, only one of two clusters (cluster 5) contained top 10 clones. The exclusive occupancy of these top 10 clones in only 5 of 6 TP1-associated clusters (clusters 0, 1, 3, 4, and 7), and in only 1 of 2 TP0-associated clusters (cluster 5) raised the possibility that a discrete, pre-existing gene expression state at TP0 confers higher drug tolerance on a subset of clones, resulting in their preferential selection by treatment (high prevalence at TP1).

**Figure 3:**
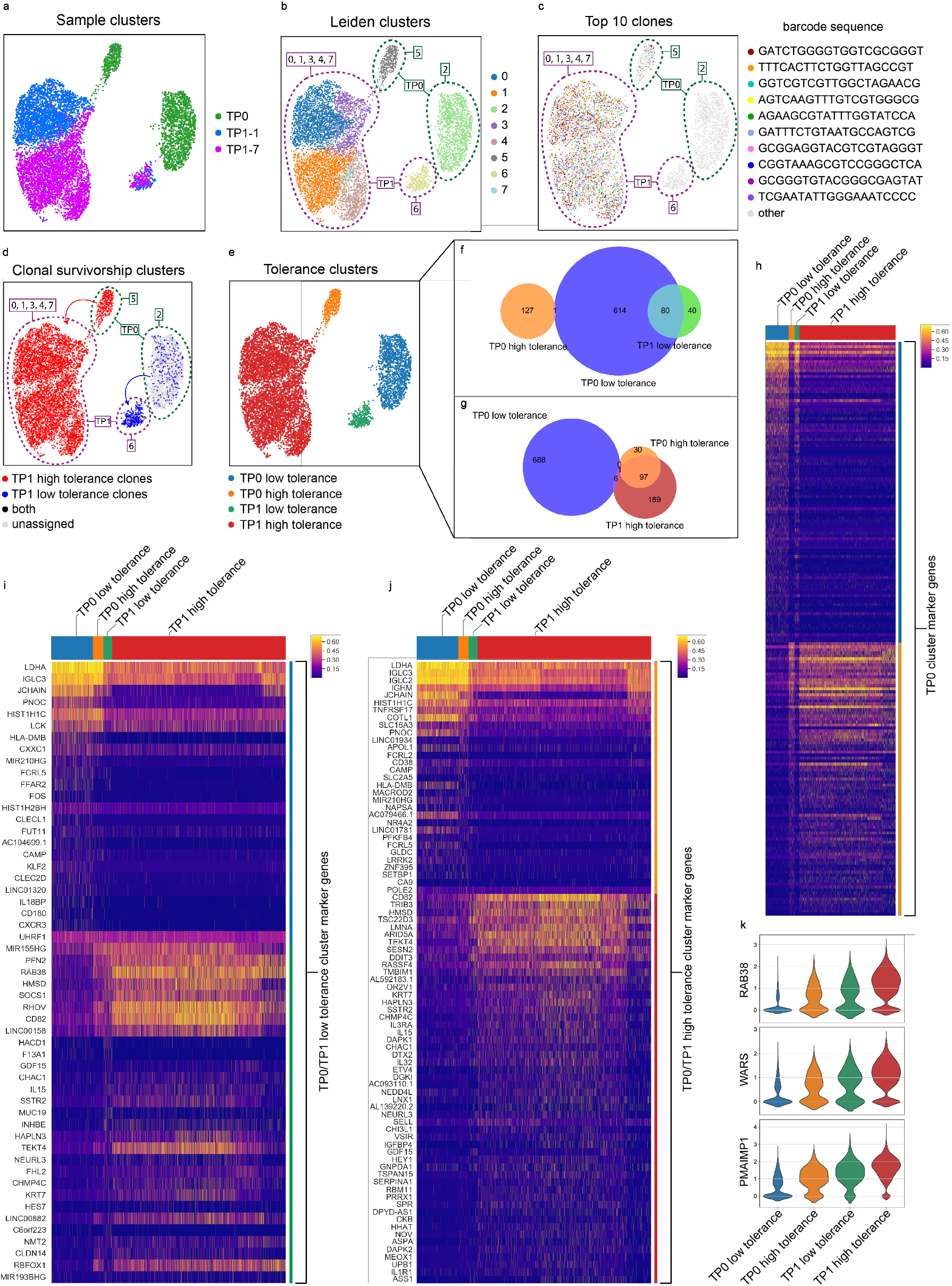
Single-cell RNA sequencing of barcoded populations prior to and following cytotoxic therapy. (a) UMAP visualization of 8,975 single-cell gene expression profiles across TP0 (n = 2,003) and TP1 replicates 1 (n = 3,134) and 7 (n = 3,838). (b) Leiden clustering reveals transcriptomic heterogeneity in TP0 and TP1 samples. Clusters 5 and 2 are exclusively comprised of cells from TP0 and clusters 0, 1, 3, 4, 6 and 7 from cells at TP1. (c) Annotation of the top 10 most abundant clones at TP1 highlights clonal relationships between TP0 and TP1 sub-populations. (d) Clones present in TP1 leiden clusters 0, 1, 3, 4 and 7 were annotated in red as ‘TP1 high tolerance clones’ (288 clones), those present in TP1 leiden cluster 6 were annotated in blue as ‘TP1 low tolerance clones’ (117 clones), and those found in both clusters were annotated in black as ‘both’ (3 clones). Projection of these clonal classifications on TP0 clusters revealed that TP1 high tolerance clones are almost exclusively present in TP0 leiden cluster 5 and TP1 low tolerance clones in TP0 leiden cluster 2. (e) UMAP cluster relabeling based off of clonal and survival relationships. (f-g) Venn diagrams demonstrating significant overlap between clones present in their respective tolerance classification pre- and post-treatment. (h-j) Heatmap of marker genes distinguishing high and low tolerance clusters at TP0 (h), upregulated in TP1 as compared to TP0 in low tolerance clusters (i), and upregulated in TP1 as compared to TP0 high tolerance clusters (j) (>2 log2 fold change, q-value <0.05). (k) Violin plots demonstrating three representative genes upregulated after treatment that were found upregulated in the TP0 high tolerance as compared to the TP0 low tolerance. Expression values in heatmaps and violin plots are log-transformed normalized counts.

To determine whether there are broader patterns of clonal survival amongst the clusters comprising TP0 and TP1, we annotated all clones present in clusters 0, 1, 3, 4, and 7 as “TP1 high tolerance clones” and those present in cluster 6 as “TP1 low tolerance clones”, with clones present in both cluster sets labeled as “both”. In this fashion, we could highlight this clonal survivorship pattern in greater detail, with clones present in TP0-associated cluster 5 clearly matching the TP1 high tolerance clones, and clones present in TP0-associated cluster 2 matching the TP1 low tolerance clones (**Fig 3d**). Specifically, we identified a total of 288 unique barcodes within the TP1 high tolerance clone classification and 117 unique barcodes within the TP1 low tolerance clone classification, with a nominal overlap of 3 barcodes shared by both subsets. Given the highly segregated patterns of gene expression, clonal composition, and survivorship, we additionally grouped and relabeled clusters 0, 1, 3, 4, and 7 as TP1 high tolerance and cluster 5 as TP0 high tolerance, as well as cluster 6 as TP1 low tolerance and cluster 2 as TP0 low tolerance (**Fig 3e**). Upon quantifying the clonal overlap between these four clusters, we observed that TP0 high tolerance and TP1 high tolerance clusters shared 97 clones, with only minimal overlap with the TP0 low tolerance cluster. As well, we found that TP0 and TP1 low tolerance clusters shared 80 clones, without any overlap with the TP0 high tolerance cluster (**Fig 3f-g**).

To understand the gene expression differences underlying these two distinct phenotypes, we performed marker gene analysis between the pre-treatment high tolerance and low tolerance populations (>0.5 log2 fold change and q-value <0.05, as described in **Methods**). These included genes in canonical CLL signaling pathways such as Wnt signaling (*WNT10A, CREM* and *TCF7)*, inflammatory/migratory *(CXCR4, PI3KR1, PI3KR5, CCL20, JAK1, RAB20, RAB27A* and *GCSAM*) and chromatin modification pathways *(PCGF5, AUTS2)* (**Table 1**). Interestingly, we also observed lower expression of type 2 antigen presentation in the high vs low tolerance population (including *CD86* and *HLA-DMA, HLA-DMB, HLA-DOA*), as well as increased expression of pro-growth and survival genes *(MAPK10, PPP3CA* and *ELK1*). Conversely, genes upregulated in the low tolerance populations included those involved in the prostaglandin biosynthesis and metabolism pathway (which have a known role in driving inflammation and migration in adjacent cells) such as *FADS1, TBXAS1* and *CBR1,* as well as those involved in the regulation of focal adhesion *(ITGB2, PAK1, FGFR1* and *VCL*). These gene expression profiles were largely maintained in their respective TP1 high and low tolerance clusters, indicating that these are, to some extent, stable phenotypes (**Fig 3h; Table 1**). Altogether, these findings indicate a role for high proliferation and migration in the high vs low tolerance population, as well as a strong anti-apoptotic phenotype in high tolerance cells.

**Table 1:**
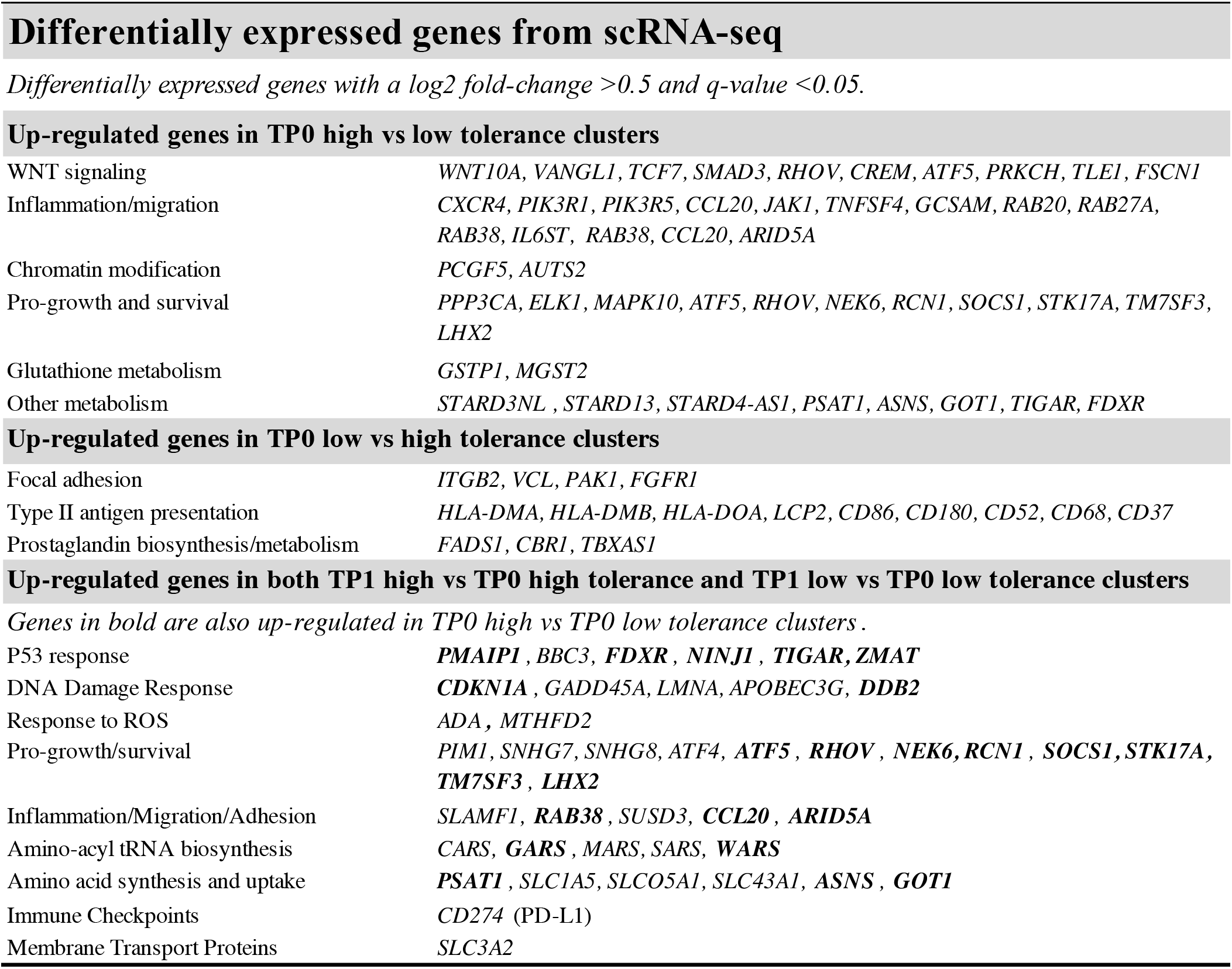
Differentially expressed genes across high and low tolerance populations. Signaling pathways and cellular mechanisms identified through differential expression analysis of gene expression clusters identified by scRNA-seq (>0.5 log2 fold change, q-value <0.05). Comparison groups include: TP0 high vs TP0 low tolerance, TP0 low vs TP0 high tolerance, FM high vs TP0 high tolerance, and FM low vs TP0 low tolerance.

A comparison of TP1 to TP0 high tolerance and TP1 to TP0 low tolerance populations revealed a number of genes that are differentially regulated by exposure to treatment (**Fig 3i-j, Table 1**). Both high and low tolerance populations experienced a strong P53 and DNA damage response *(PMAIP1, BBC3, CDKN1A, GADD45A, LMNA)* as well as upregulation of genes involved in response to reactive oxygen species (ROS) (ADA and *MTHFD2*). Both subpopulations also appeared to experience upregulation of proteins involved in aminoacyl t-RNA biosynthesis, which have previously been described in fludarabine-resistant lymphomas and in chemotherapeutic resistance in other cancer types (Lorkova et al., 2015). Of note, many genes involved in these strong drug-response pathways already had high basal expression in the TP0 high vs low tolerance population (including *PMAIP1, FDXR, NINJ1, TIGAR, ZMAT, DDB2, RHOV, NEK6, RCN1, SOCS1, STK17A, TM7SF3, LHX2, RAB38, CCL20, ARID5A, GARS, WARS, PSAT1, ASNS* and *GOT1*), indicating potential priming in its response to therapy (**Fig 3k; Table 1**).

## Discussion

Recent efforts have increasingly focused on understanding the evolutionary dynamics that lead to drug resistance in cancer. Numerous genome sequencing studies have inferred tumor evolution over the course of therapy by measuring changes in SNP/allele frequencies, but have had limited success in resolving these complex phenomena in granular clonal detail (Ding et al., 2013; Jolly and Van Loo, 2018; Landau et al., 2015; McGranahan et al., 2015; Shin et al., 2017; Turajlic et al., 2015). To directly measure clonal growth dynamics, new approaches have utilized highly-sensitive lineage tracing techniques such as DNA barcoding (Blundell and Levy, 2014; Kebschull and Zador, 2018). While DNA barcoding has provided the resolution to uncover evidence of pre-existing and *de novo* alterations leading to therapeutic evasion, this approach has a very limited capacity to uncover the genomic or transcriptomic content of clones within a population (Bhang et al., 2015; Hata et al., 2016).

In this study we address these measurement limitations by implementing a combined approach, integrating DNA barcoding with a specialized sgRNA expression/capture vector, CROP-seq, to study the clonal response of the HG3 CLL model cell-line to cytotoxic therapy of fludarabine and mafosfamide (Datlinger et al., 2017). Together, these approaches have enabled the a) tagging of >1 million HG3 clones, b) measurement of clonal frequency at depths greater than 1:1×10^6^ cells, and c) clonally-resolved single-cell gene expression analysis in response to therapy. Further, the CROP-seq expressed barcode system is compatible with COLBERT clonal isolation such that clones of interest can be identified and isolated from the population for future live-cell or endpoint molecular analysis (Al’Khafaji et al., 2018).

Previous bulk WES studies in CLL have identified that drug-resistant populations can arise from small, pre-existing populations (Landau et al., 2013). Here, we similarly report the existence of a small subpopulation of HG3 cells that are highly tolerant to combined fludarabine/mafosfamide treatment, with the added resolution of direct clonal measurements using DNA barcoding and single-cell transcriptomics. First, through barcode sequencing we observed that clonal diversity was reduced by ~80% in each of 8 parallel replicates following fludarabine/mafosfamide treatment, and that a small number of clones are consistently and strongly selected for across replicates. Approximately 94% of surviving cells in each replicate had a clonal identity that was present in all 8 replicates, indicating the likelihood of a shared mode of resistance that was already pre-existing in the population.

Second, through clonally-resolved scRNA-seq of the pre-treatment and select post-treatment samples, we determined the underlying clonal composition of this HG3 population and observed that the constituent clones exhibit stable and discrete gene expression states that differentially respond to chemotherapy. Specifically, we identified two distinct gene expression clusters prior to treatment that each served as clonal reservoirs for future surviving cells. The first cluster (TP0 high tolerance) is a minor subset (~20% of TP0) that almost exclusively consists of clones that go on to survive treatment (~95% of TP1). The second cluster (TP0 low tolerance) consists predominantly of cells that are sensitive to treatment, however a small number of clones in this cluster managed to survive TP1 treatment (and go on to comprise 5% of TP1). To characterize the cellular mechanisms underlying the high and low tolerance phenotypes, we performed pathway and gene set enrichment analysis of the differentially expressed genes between TP0 high and low tolerance clusters and found that the high tolerance cluster is characterized by a marked upregulation of common CLL signaling pathways such as WNT and CXCR4, along with an inflammatory/migratory phenotype and a reliance on chromatin modification pathways. The upregulation of these key CLL pathways in the high tolerance population is of particular interest given that *FAT1* (WNT regulator) mutations and CXCR4 upregulation are frequently seen in chemo-refractory CLL (Burger and Kipps, 2006; Messina et al., 2014). The TP0 low tolerance, on the other hand, exhibited upregulated type 2 antigen presentation and prostaglandin biosynthesis/metabolism, which has a known role in driving inflammation and migration in adjacent cells (Wang and DuBois, 2006). Interestingly, many of the genes upregulated at TP1 were already upregulated in the high tolerance clusters at TP0.

Unlike targeted therapies, which usually have more specific mechanisms of resistance directly related to the pathways impacted by the targeted agent, chemotherapeutics can have more broad and varied mechanisms of resistance (Lorkova et al., 2015). Resistance to fludarabine, a purine analog, has largely involved genes involved in the nucleotide salvage pathways, anti-apoptotic pathways, transcription factors and mediators of genotoxic stress (Lorkova et al., 2015), while resistance to cyclophosphamide, an alkylating agent, has been tied to upregulated DNA repair processes and detoxification of cyclophosphamide and its metabolites (Ludeman and Gamcsik, 2002). Consistent with resistance mechanisms described in the literature, we observed an upregulation of genes involved in P53 and DNA damage response, proliferation/survival, inflammation and migration, amino-acyl tRNA synthesis and membrane transport proteins across all TP1 populations. Of note, the majority of genes involved in these drug-response pathways were already upregulated in the high tolerance population prior to treatment, perhaps contributing to the increased survival of these clones.

In sum, the ability to leverage these multifunctional lineage tracing approaches enables an unprecedented characterization of complex systems. By tagging large heterogenous cancer populations and tracking clonal abundancies as well as clonally resolved single-cell gene expression profiles over time, we can measure the range of clonal responses to environmental stressors. Further, with this combined approach, we can also leverage COLBERT to derive clones of interest and isolate clones for live-cell characterization. Herein, we demonstrate one application of how these approaches can come together to uncover the clonal survival and gene-expression dynamics that underlie cellular responses to cytotoxic therapy. Knowledge of these evolutionary paths will contribute to our collective understanding of how therapy can shape tumors over time and highlight new areas of therapeutic opportunity.

## Methods

### Cell culture

The HG3 cell line (DSMZ #ACC 765) was cultured in RPMI 1640 (Gibco #11875-093) with 15% fetal bovine serum (FBS) (Sigma-Aldrich), 1% GlutaMAX (Gibco, cat# 35050-061) and 1% penicillin-streptomycin (Life Technologies). Cells were incubated at 37 °C, 5% CO2 and passaged every 48 hours. All subsets of the HG3 cell line were also cultured under these conditions. Cell line identity was confirmed through fluorescence in situ hybridization demonstrating the characteristic *del*(13q). Cells were routinely tested for mycoplasma per the manufacturer’s instructions (VenorGeM Mycoplasma Detection Kit; Sigma-Aldrich #MP0025).

### CROP-seq-BFP-TSO-Guide vector assembly

The CROP-seq-Guide-Puro vector (Addgene, #86708) was modified by replacing the puromycin resistance gene with BFP. The restriction enzymes PspLI (Thermo Fisher #FERFD0854) and MluI (Thermo Fisher #FERFD0564) were used to digest and remove the puromycin resistance marker and a gBlock^®^ of the BFP fluorescent marker was cloned into the digested vector using the same restriction site overhangs. To increase transcript capture during RNA sequencing workflows, the template switch oligo sequence (AAGCAGTGGTATCAACGCAGAGTACATGGG) was cloned upstream of the hU6 promoter following digestion of the CROP-seq-Guide-BFP vector with KflI (Thermo Fisher FERFD2164), generating the new vector ‘CROP-seq-BFP-TSO-Guide’.

### High-Complexity Barcode-sgRNA Library Construction

A 60 bp oligonucleotide containing a 20 bp random sequence corresponding to the barcode-sgRNA (BgL-BsmBI) and a reverse extension primer (RevExt-BgL-BsmBI) were ordered from IDT:

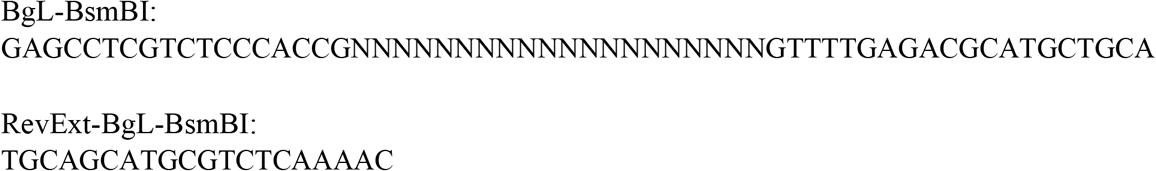

The following extension reaction was performed to generate the double-stranded barcode-sgRNA oligo: 10 μL Q5 PCR Reaction Buffer, 1 μL 10 mM dNTP mix, 1 μL of 100 μM BgL-BsmBI barcoded template, 2 μL of 100 μM RevExt-BgL-BsmBI reverse extension primer, 0.5 μL Q5 polymerase, and 35.5 μL water and incubated as follows: 98°C for 2 min, 10x (65°C for 30 s, 72°C for 10s), 72°C for 2 min, hold at 4°C. The double stranded barcode-sgRNA oligo was purified using a QIAquick PCR Purification Kit (Qiagen, #28104). The double-stranded product contains two BsmBI sites that, upon digestion, generate complimentary overhangs for ligation into CROP-seq-BFP-TSO-Guide. 1.5μg of BsmBI digested (Thermo Fisher #FERFD0454) CROP-seq-BFP-TSO-Guide vector was entered into a Golden Gate assembly reaction with the double-stranded barcode-sgRNA insert at a molar ratio 1:5. The reaction was incubated at 100x (42°C for 2 min and 16°C for 5 min). This reaction was purified and concentrated in 12 μl water using the DNA Clean & Concentrator™ kit (Zymo, #D4033) and transformed into electrocompetent SURE 2 cells (Agilent, #200152). Transformants were inoculated into 500 ml of 2xYT media containing 100 μg/ml carbenicillin and incubated overnight at 37°C. Bacterial cells were pelleted by centrifugation at 6,000 RCF at 4°C for 15 minutes and plasmid DNA was extracted using a Qiagen Plasmid Plus Midi Kit (Qiagen #12943). To estimate the total number of unique barcodes present in the plasmid pool, we used a fourth-order polynomial regression to fit the sampled data. This function plateaus at approximately 7.6×10^6^ barcodes, corresponding to the projected number of unique barcodes present in the plasmid pool.

### Lentivirus production of barcode-sgRNA libraries

Twenty-four hours prior to lentiviral transfection, HEK293T cells were plated in a 6-well plate at 600,000 cells per well in 3 mL of antibiotic-free DMEM with 10% FBS. When cells reached 70-80% confluency, they were transfected with 9 μL of Lipofectamine 2000 (Thermo Fisher #11668027), 1.5 μg per well of PsPax2 (Addgene #12260), 0.4 μg/well of VSV-G (Addgene #8454), and 2.5 μg of CROP-seq-BFP-TSO-Barcode_sgRNA. Twenty-four hours post-transfection, media was exchanged for 3 mL of antibiotic-free DMEM with 20% FBS. Media containing viral particles was collected at 48 and 72 hours posttransfection, pooled, filtered through a 0.45 μm Nalgene syringe filter (Thermo Scientific #723-2945) coupled to a 10 mL syringe, and subsequently concentrated using a 50 mL size-exclusion column (Vivaspin20 30,000 MWCO PES from Sartorius, #VS2022) by centrifugation at 2,200 RCF at 4°C for 2 hours. Concentrated virus was then removed from the upper chamber and stored at −80°C in 50 μL aliquots for later use.

### Lentiviral transduction with barcode-sgRNA libraries

Prior to transduction with the lentiviral barcode library, HG3 cells were seeded in a 6-well plate at 2×10^6^ cells per well in 3 mL of culture media. Cells were transduced with the barcoded CROP-seq-BFP-TSO-Barcode_sgRNA lentivirus using 0.8 μg/mL of polybrene and were subsequently centrifuged at 500 RCF for 1.5 hours at 37°C, followed by overnight incubation at 37°C, 5% CO2. To reduce the likelihood that two viral particles enter a single cell, the MOI was kept below 0.1. Sixteen hours after incubation, the transduction media was exchanged for fresh culture media. After 48 hours, 1.2×10^6^ BFP+ cells were isolated by FACS and subsequently cultured for 27 days to 2.4×10^8^ cells, after which they were cryopreserved in 12 vials of 2×10^7^ cells each.

### Chemotherapy-resistance studies in HG3 cells

Dose response curves were established by plating 200,000 cells in 200 μL and treating with 0.1 μM, 0.5 μM, 1 μM, 5 μM, 10 μM and 50 μM of either fludarabine (Selleckem, #S1491), mafosfamide (Santa Cruz, #SC-211761), or fludarabine and mafosfamide combined. After 72 hours, viability was determined using CellTiter-Glo (Promega, #G7570). A lethal dose of 95% reduction in viability (5 μM fludarabine + 5 μM mafosfamide) was chosen to ensure stringent selection of the barcoded population. Prior to drug treatment, a cryovial containing 1.5×10^7^ viable barcoded HG3 cells was thawed and expanded for 8 days, to a cell count of 2.4×10^8^. Barcoded cells were then plated in 8 parallel T25 flasks, each at a density of 1×10^7^ cells in 10 mL of culture media. Cells were immediately treated with an LD95 dose of fludarabine + mafosfamide and incubated in drug media for 72 hours, at which point cells were washed twice with PBS and resuspended in 10 mL of fresh media and plated in 8 new T25 flasks. Upon repopulation of the flask (21 days post-treatment), 2 mL of cells and media (1/5 of the flask) were pelleted for genomic DNA extraction. The remaining cells were cryopreserved in FBS + 10% DMSO for future analysis.

### Next-generation barcode sequencing

Genomic DNA was extracted from pre- and post-treated cell populations using the DNeasy Blood & Tissue Kit (Qiagen #69504) per the manufacturer’s instructions. Barcode sequences were amplified by PCR using primers containing flanking barcode annealing regions and Illumina adapter/6-bp index sequences. To sequence the barcoded CROP-seq-BFP-TSO-Barcode_sgRNA, 50 ng of plasmid DNA were used as a template. To sequence the barcoded populations, 2 μg of genomic DNA were used per PCR reaction in 8 parallel PCR reactions. In total, 36 μg of genomic DNA were used for TP0 and 16 μg of genomic DNA for TP1 replicates. All reactions were purified using a 1.8x AMPureXP bead cleanup (Beckman Coulter #A63880) and sequenced on an Illumina NextSeq platform by single-end sequencing for 75 cycles.

### Barcode Sequencing Analysis

Raw barcode sequencing data from barcode gDNA amplifications was processed with cutadapt (https://cutadapt.readthedocs.io/en/stable/) to identify reads containing barcodes and trim the 25 bp flanking adapter sequences (5’: ATCTTGTGGAAAGGACGAAACACCG, 3’: GTTTTAGAGCTAGAAATAGCAAGTT). To accommodate barcode reads with sequencing errors, flanking sequence identification was permitted a maximum expected error of 0.1. Positively identified and trimmed barcode sequences 20 bp in length and with no ambiguous (N) bases were included in further analyses. Barcode sequences were then filtered for read quality (requiring all bases to have a minimum PHRED of 20). To compensate for amplification artifacts and sequencing error, trimmed barcode sequences were tightly clustered across samples with the Starcode tool (https://github.com/gui11aume/starcode) using a message-passing algorithm with a maximum Levenshtein distance of 1 and minimum radius size of 3 (Zorita et al., 2015). Remaining singleton barcodes that were not clustered were removed, resulting in a median of 1.13×10^7^ barcode sequences per post-treatment sample (min 9.7×10^6^, max 1.35×10^7^) and 2.06×10^8^ sequences for the TP0 sample.

### Single-cell RNA sequencing

Cryopreserved samples from TP0 and TP1 Replicates 1 and 7 were thawed, pelleted and resuspended in cold PBS-0.04% BSA. 100,000 viable BFP+ cells were sorted into cold PBS-0.04% BSA, resuspended at 1×10^6^ cells per mL, and barcoded with a 10X Chromium Controller (10X Genomics) by reverse transcription of the RNA in individual cells. Sequencing libraries were subsequently constructed using reagents from a Chromium Single Cell 3’ v2 reagent kit according to the manufacturer’s instructions (10X Genomics, #PN-120267). Sequencing was performed on an Illumina NovaSeq platform (paired-end, Read 1: 26 cycles and Read 2: 98 cycles) according to the manufacturer’s instructions (Illumina).

### Single-cell RNA sequencing Analysis

#### Raw Sequence Processing and Alignment

Using Cell Ranger (v3.0) (10XGenomics), the aggr command was used to aggregate the three libraries (with the default parameters) to generate a single gene-count matrix.

#### Filtering and Normalization

The filtered matrices produced by Cell Ranger were loaded into scanpy (v1.4.3) (Wolf et al., 2018). Cells were annotated by sample and clonal membership. Only cells meeting the following requirements were retained for further analysis: (a) a minimum of 4000 transcript counts, (b) a maximum of 8% of counts attributed to mitochondrial genes, and (c) a minimum of 750 genes detected. Normalization was conducted based on recommendations from multiple studies that compared several normalization techniques (Büttner et al., 2019; Luecken and Theis, 2019; Vieth et al., 2019). Data normalization was performed by implementing the following: preliminary clustering of cells by constructing a nearest network graph and using scanpy’s implementation of Leiden community detection (Traag et al., 2019), calculating size factors using the R package scran (v1.10.2) (Lun et al., 2016), and dividing counts by the respective size factor assigned to each cell. Normalized counts were then transformed by adding a pseudocount of 1 and taking the natural log.

#### Regressing Out Cell Cycle Expression Signatures

Using a list of genes known to be associated with different cell cycle phases (Tirosh et al., 2016), cells were assigned S-phase and G2M-phase scores. This dataset was then run through scanpy’s *regress_out* function with both score categories as inputs.

#### Assigning Cells to Transcriptomically-Distinct Clusters

Every gene was centered to a mean of zero and principal component analysis was performed. The top 40 principal components were used to construct a nearest neighbor graph. Cells were then assigned to clusters using the Leiden algorithm with the resolution set to 0.65, resulting in 8 clusters.

#### Differential Expression Analysis

Differential expression for each gene was assessed using scanpy’s *rank_genes_groups* function with Wilcoxon tests to generate p-values. A truncated normal (TN) test (Zhang et al., 2019) was also performed on all genes, yielding a second set of p-values. Each set of p-values was adjusted using Benjamani-Hochberg false discovery rate correction, producing q-values. To be conservative, “consensus” q-values were calculated by selecting the larger of the two q-values for each gene. A gene was considered statistically significant if its consensus q-value was <0.05. Pathways analysis was performed using the MetaCore software (https://portal.genego.com/cgi/data_manager.cgi). Gene lists for Table 1 were compiled from top scoring pathways.

### Clonal Assignment in scRNA-seq Data

Cell- and UMI-labeled sequencing data in the unaligned BAM file reads from the 10X Cell Ranger platform were used to identify clone barcodes for each cell. The cutadapt tool was used to identify adapters flanking the barcode sequences of interest (5’: ATCTTGTGGAAAGGACGAAACACCG, 3’: GTTTTAGAGCTAGAAATAGCAAGTT) – however, due to the extreme 3’ nature of the 10X data, only the 3’ adapter was required for barcode identification and a minimum of 10 bp of the barcode portion of the read were required. To correct for processed barcode reads less than 20 bp, all barcode reads within a particular cell were compared. If an exact match to the 3’ end of a longer barcode read was found, the read was extended with the matching sequence. Length-corrected clone barcode reads were omitted if not found in the whitelist of known clone barcodes from the original TP0 population. For each cell, UMI-replicate reads were resolved by taking the mode barcode read for each UMI. To resolve instances where multiple clone reads in a particular cell were observed, we annotated cells by their maximum clone barcode given no other clone barcode surpassing 50% of the read count of the maximum clone. Clonal reads from cells with many major clone barcodes that could not be resolved were removed.

## Acknowledgements

We acknowledge support from the National Institutes of Health (5R21CA212928, 5R01CA226258 to A.B.; 3P01CA206978-03S1, 1U10CA180861-01, 1P01CA206978-01 to C.J.W.). C.J.W. is a Scholar of the Leukemia and Lymphoma Society and C.G. is a Scholar through the American Society of Hematology MMSAP Program and the F31 Diversity Individual Predoctoral Fellowship program through the NCI. E. B. is a recipient of the University of Texas at Austin Provost’s Graduate Excellence Fellowship and the F. M. Jones & H.L. Bruce Endowed Graduate Fellowship. K.J. is grateful for support through an NSF Graduate Research Fellowship (1610403). We thank the Dana-Farber Flow Cytometry Core, the Broad Institute Walk-Up Sequencing Core, and the Genome Sequencing and Analysis Facility at the University of Texas at Austin for their services. All scRNA-seq workflows were conducted in the Translational Immunogenomics Lab at DFCI.

## Author Contributions

A.A. and C.G. designed the research and performed the bulk of the experiments, under supervision by A.B. and C.J.W. S.L. and K.J.L. prepared the 10X Genomics scRNA-seq libraries. E.B. and R.D. generated the computational pipelines. E.B. and D.N. contributed to statistical analyses. K.E.J. and W.Z. contributed to data analysis and validation. All authors participated in data analysis. A.A. and C.G. wrote the manuscript. All authors reviewed the manuscript prior to submission.

